# Genomic bases of insularity and ecological divergence in barn owls (*Tyto alba*) of the Canary Islands

**DOI:** 10.1101/2021.12.24.473866

**Authors:** Tristan Cumer, Ana Paula Machado, Felipe Siverio, Sidi Imad Cherkaoui, Inês Roque, Rui Lourenço, Motti Charter, Alexandre Roulin, Jérôme Goudet

**Affiliations:** Department of Ecology and Evolution, University of Lausanne, Lausanne, Switzerland; Canary Islands’ Ornithology and Natural History Group (GOHNIC), 38480 Buenavista del Norte, Tenerife, Canary Islands, Spain; Ecole Supérieure de Technologie de Kénitra, Ibn Tofail University, Kénitra, Morocco; Mediterranean Institute for Agriculture, Environment and Development, Laboratory of Ornithology, IIFA, University of Évora, Évora, Portugal; Shamir Research Institute, University of Haifa, Katzrin, Israel; Department of Geography and Environmental Sciences, University of Haifa, Haifa, Israel; Swiss Institute of Bioinformatics, Lausanne, Switzerland

**Keywords:** Local adaptation, Niche analysis, Population genomics, *Tyto alba gracilirostris*, Whole genome sequencing

## Abstract

Islands, and the particular organisms that populate them, have long fascinated biologists. Due to their isolation, islands offer unique opportunities to study the effect of neutral and adaptive mechanisms in determining genomic and phenotypical divergence. In the Canary Islands, an archipelago rich in endemics, the barn owl (*Tyto alba*) is thought to have diverged into a subspecies (*T. a. gracilirostris*) on the eastern islands, Fuerteventura and Lanzarote. Taking advantage of 40 whole-genomes and modern population genomics tools, we provide the first look at the origin and genetic makeup of barn owls of this archipelago. We show that the Canaries hold diverse, long-standing and monophyletic populations with a neat distinction of gene pools from the different islands. Using new method, less sensitive to structure than classical *F*_ST_, to detect regions involved in local adaptation to the insular environment, we identified a haplotype-like region likely under positive selection in all Canaries individuals. Genes in this region suggest morphological adaptations to insularity. In the eastern islands, where the subspecies *T. a. gracilirostris* is present, genomic traces of selection pinpoint signs of locally adapted body proportions and blood pressure, consistent with the smaller size of this population living in a hot arid climate. In turn, genomic regions under selection in the western barn owls from Tenerife showed an enrichment in genes linked to hypoxia, a potential response to inhabiting a small island with a marked altitudinal gradient. Our results illustrate the interplay of neutral and adaptive forces in shaping divergence and early onset speciation.

## Introduction

Due to the often-peculiar organisms that inhabit them, islands have always fascinated naturalists and scientists alike. Since Darwin’s first visit to the Galapagos in 1835, the study of insular populations has been crucial to the development of evolutionary theory (MacArthur and Wilson 1967; Grant 1998; Warren et al. 2015). Labelled nature’s test tubes, islands are home to a myriad of endemic (sub)species. Moreover, the combination of their relatively small size, discrete borders, geographical isolation and natural replication provides an excellent setting to study the evolutionary forces underlying population divergence and speciation (Losos and Ricklefs 2009). The divergence of insular populations from their founders and surrounding islands is the result of neutral and selective forces. Disentangling the respective impacts of these two forces is a challenging task, as they are interconnected and can both contribute to genetic and phenotypic differentiation.

Isolated and small populations, such as those frequently found on islands, are under a markedly strong influence of genetic drift. This process will alter the genetic makeup of the population by randomly removing rare alleles and fixating common ones, hence decreasing genetic diversity (Frankham 1997). This is a common occurrence on islands which, coupled with low gene flow, can lead to inbreeding (Keller and Waller 2002) and accelerate neutral divergence. Conversely, the absence of regular gene flow, and the often distinctive ecological conditions of the islands, can facilitate the action of local adaptation on beneficial alleles (Lenormand 2002; Tigano and Friesen 2016). This process can lead to the emergence of ecomorphs via ecological divergence as populations adapt to unfilled insular niches (Losos and Ricklefs 2009), particularly so in remote islands that are colonized less often (MacArthur and Wilson 1963). Ecomorphs can occur in different islands or in the same one (Gillespie et al. 1997; Losos et al. 1998; Gillespie 2004), a concept mirrored in inland lakes (Malinsky et al. 2015), the aquatic homologous of islands. Eventually, ecomorphs can become new subspecies or even species and, in extreme cases, result in adaptive radiations as illustrated by Darwin’s finches (Grant 1999; Lamichhaney et al. 2015).

The Afro-European barn owl (*Tyto alba*) is a non-migratory, nocturnal raptor present from Scandinavia to Southern Africa. It is also found on numerous islands and archipelagos where subspecies have often been described (Uva et al. 2018). A recent study of the species’ genetic structure in the Western Palearctic (Cumer et al. 2021) described two main lineages occupying this region: the eastern lineage in the Levant and the western in Europe. In addition, Cumer et al. (2021) showed that barn owls from Tenerife (Canary Islands) were very distinct from both lineages. The Canaries are a volcanic archipelago that was formed several million years ago (Anguita and Hernán 2000) about 100 km from the coast of north-western Africa and was never connected to the mainland. This long-term isolation (Norder et al. 2019), along with its subtropical climate and elevation gradients (Steinbauer et al. 2016), has resulted in a high level of endemicity, for example in plants (Carine et al. 2009), reptiles (Thorpe and Baez 1993; Nogales et al. 1998; Nogales et al. 2001; Molina-Borja 2003; Mateo et al. 2011), mammals (Hutterer et al. 1987; Pestano et al. 2003; Firmat et al. 2010; Masseti 2010) and birds (Illera et al. 2016; Lifjeld et al. 2016; Rodríguez et al. 2020; Senfeld et al. 2020). Among them, an endemic barn owl subspecies, *T. a. gracilirostris* (Hartert, E, 1905), has been described based on its morphological traits, especially for its smaller size and darker coloration (Bannerman 1963; Clements et al. 2019). This subspecies is recorded in the eastern Canary Islands of Lanzarote and Fuerteventura, as well as in its surrounding islets (Lobos, Alegranza, Montaña Clara and La Graciosa), and is the only barn owl in this sector of the archipelago (Siverio 2007).

The presence of this subspecies on the eastern islands is surprising however, given that it is sandwiched between the western islands and the mainland which both harbour the nominal species *T. alba.* Lacking any evidence of different colonization origins or timing, this could suggest that selection to local conditions acting on the eastern population has accelerated divergence in comparison to the western one. In both cases, neutral and adaptive microevolutionary processes promoting divergence of insular populations would leave traces on their genomic makeup. The advances in sequencing technology, and its decreasing costs, now allow to sequence the entire genomes of individuals. With the parallel development of sophisticated tools, it is possible to analyse changes in allelic frequencies at a high resolution to investigate the history of populations and inspect the genomic landscape for signals of local adaptation.

Here, we investigate the genomic bases of differentiation of barn owls from the Canary Islands. Making use of the whole-genome sequences from 40 individuals, we first describe the neutral genetic structure and diversity of the Canaries populations in contrast to the mainland, in order to retrace their history. Second, we employ a new relatedness-based method to probe the genomic landscape of these isolated populations for signals of local adaptation to the insular environment in regards to the mainland. Third, we characterize the climatic niches barn owls occupy on eastern and western islands, and explore how each population is diverging to adapt to their niches from a genomic perspective. Our results elucidate the genomic bases of the differentiation of insular populations, thus enlightening the classification of the Canaries barn owls.

## Materials & Methods

### Whole-genome sequencing, SNP calling and identification of coding regions

For this study, 40 individual barn owls were sampled from four populations (Figure 1; Supplementary Table 1): Eastern Canaries (EC, Fuerteventura and Lanzarote), Western Canary (WC, Tenerife), Morocco (MA), Portugal (PT) and Israel (IS). All but the EC and MA populations were published in previous work, including a North American owl that was used as an outgroup for some analyses (Genbank accession JAEUGV000000000 (Cumer et al. 2021; Machado et al. 2021). For EC and MA, we followed the same molecular and sequencing protocol as in the aforementioned publications. Succinctly, genomic DNA was extracted using the DNeasy Blood & Tissue kit (Qiagen, Hilden, Germany) and individually tagged 100bp TruSeq DNA PCR-free libraries (Illumina) were prepared according to manufacturer’s instructions. Whole-genome resequencing was performed on multiplexed libraries with Illumina HiSeq 2500 high-throughput paired-end sequencing technologies at the Lausanne Genomic Technologies Facility (GTF, University of Lausanne, Switzerland).

**Figure 1.**
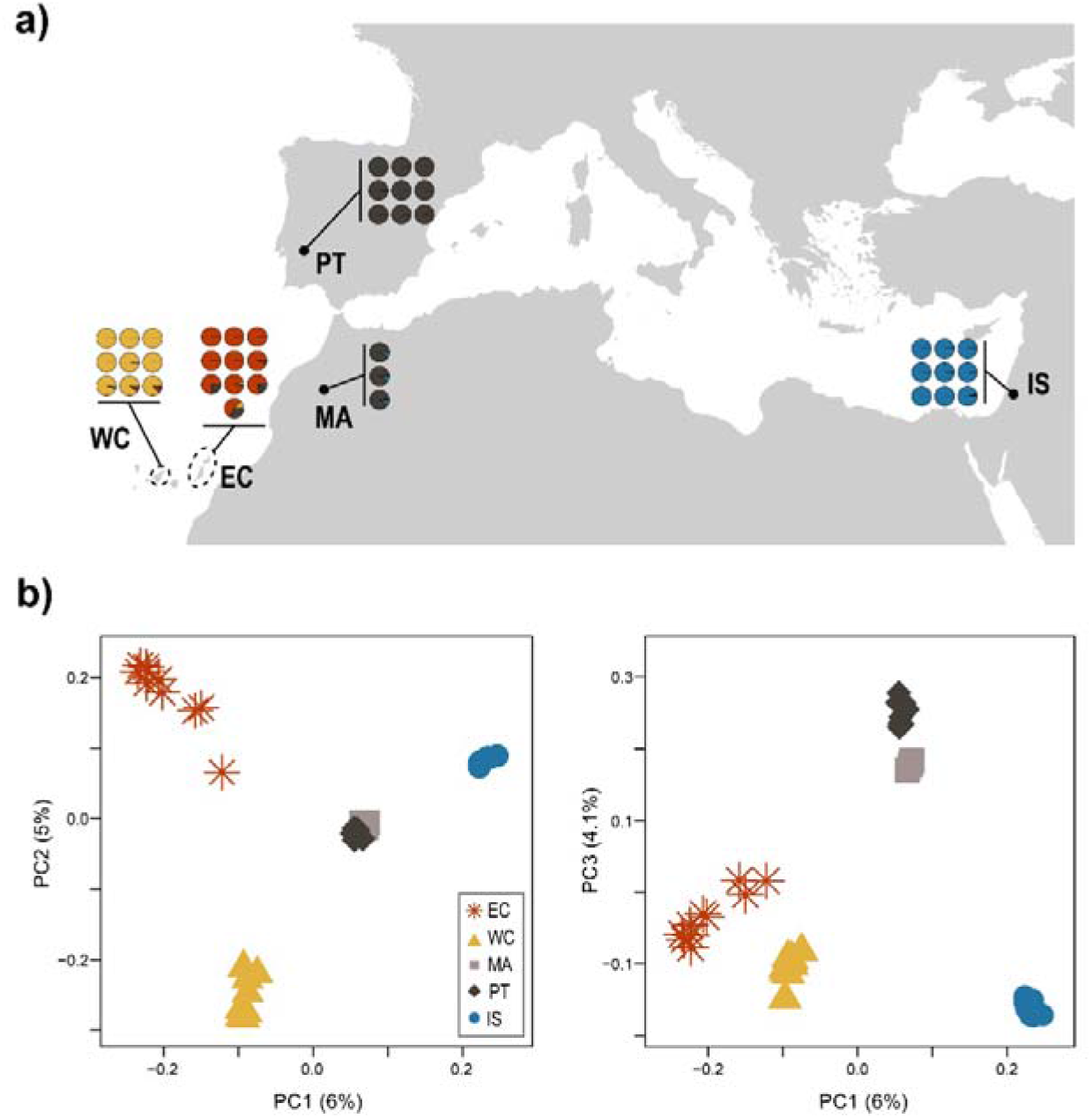
Population structure of barn owls from the Mediterranean Basin and the Canary Islands. **a)** Individual admixture proportion of each of K=4 lineages as determined by sNMF. Black dots are located at the approximate centroid of each sampled population. Dashed lines encircle the island(s) sampled for each Canary population. **b)** PCA of the 40 individuals. Point shape and colour denote populations according to the legend. Axes one to three are shown and values in parenthesis indicate the percentage of variance explained by each axis.

The bioinformatics pipeline used to obtain analysis-ready SNPs from the 40 individuals plus the outgroup was the same as in (Machado et al. 2021), adapted from the Genome Analysis Toolkit (GATK) Best Practices (van der Auwera et al. 2013) to a non-model organism following the developers’ recommendations. Briefly, reads were trimmed with Trimommatic v.0.36 (Bolger et al. 2014) and aligned with BWA-MEM v.0.7.15 (Li and Durbin 2009) to the barn owl reference genome (GenBank accession JAEUGV000000000; Machado et al. 2021). Base quality score recalibration (BQSR) was performed in GATK v.4.1.3 using high-confidence calls obtained from two independent callers: GATK’s HaplotypeCaller and GenotypeGVCF v.4.1.3 and ANGSD v.0.921 (Korneliussen et al. 2014). Following BQSR, variants were called with GATK’s HaplotypeCaller and GenotypeGVCFs v.4.1.3 from the recalibrated bam files. Genotype calls were filtered using GATK and VCFtools (Danecek et al. 2011) if they presented: low individual quality per depth (QD < 5), extreme coverage (600 > DP > 1000), mapping quality (MQ < 40 and MQ > 70), extreme hetero or homozygosity (ExcessHet > 20 and InbreedingCoeff > 0.9) and high read strand bias (FS > 60 and SOR > 3). We filtered further at the level of individual genotype for low quality (GQ < 20) and extreme coverage (GenDP < 10 and GenDP > 40). Lastly, we kept only bi-allelic sites with less than 5% of missing data across individuals resulting in 7′283′516 SNP. For analyses of neutral population structure and demography, an exact Hardy-Weinberg test was used to remove sites that significantly departed (p<0.05) from the expected equilibrium using the R (R Development Core Team 2016) package HardyWeinberg (Graffelman and Morales-Camarena 2008; Graffelman 2015) yielding 6’827’220 SNP with a mean coverage of 19.9X (4.6 SD).

### Population structure and genetic diversity

To investigate population structure among our samples, sNMF v.1.2 (Frichot et al. 2014) was run for K 2 to 5 in 25 replicates to infer individual clustering and admixture proportions. For this analysis, singletons were excluded and the remaining SNPs were pruned for linkage disequilibrium in PLINK v1.946 (Purcell et al., 2007; parameters-indep-pairwise 50 10 0.1) as recommended by the authors, retaining 288′775 SNP. The same dataset was used to perform a Principal Component Analysis (PCA) with the R package SNPRelate (Zheng et al. 2012). A second PCA was performed by merging the data in this study with that of (Cumer et al. 2021) that includes European populations (total of 2’036’320 SNP with no missing data) to assess where the Eastern Canary population (unsampled in Cumer et al. 2021) falls among the larger sampling.

Individual observed heterozygosity and population-specific private alleles were estimated using custom R scripts for each population. Individual-based relatedness (β; (Weir and Goudet 2017; Goudet et al. 2018), inbreeding coefficients for whole genome SNP data (*F*_IS_ and *F*_IT_), overall, population specific and pairwise *F*_ST_ (Weir and Goudet 2017) were calculated with hierfstat v.0.5-9 (Goudet 2005).

### Demographic history

To investigate the demographic history of the insular populations and potential admixture events we used Treemix (Pickrell and Pritchard 2012). Using the LD-pruned dataset filtered further to include no missing data (228’980 SNP), Treemix was run for 10 replicates with 0 to 5 migration events, rooting the tree on the IS population, given its position on the PCA and what is known of the region from previous work (Cumer et al. 2021). The function get_f was used to estimate the variance explained by adding migration events.

### Detection of genomic regions under selection

#### Insular vs mainland barn owls

In this study, we aimed to identify genomic regions potentially under selection at two different levels. First, to detect signatures of selection specific to barn owls of the Canary Islands, we grouped insular individuals (EC and WC) and compared them to the mainland ones (MA, PT and IS). A script by Simon Martin (https://github.com/simonhmartin/genomics_general/blob/master/popgenWindows.py) was used to estimate genome wide patterns of relative (*F*_ST_^ILvsML^) and absolute (*d*_xy_) divergence between the insular and mainland groups, and to calculate nucleotide diversity (π) per group, in windows of 100kbp with 20kbp steps.

SNPrelate was used to calculate a pairwise matrix of linkage disequilibrium (*r*) from SNPs with over 5% minor allelic frequency (MAF) over all insular individuals (EC and WC) on one hand and all mainland individuals (PT, MA and IS) in the other, which was then squared to obtain *r*^2^. For plotting, we estimated the mean of non-overlapping 100 SNP windows.

The topology weighting method implemented in Twisst (Martin and Van Belleghem 2017), was used to quantify the relationships between the five populations in our dataset and visualize how they change along the genome. Twisst estimated the topology based on trees produced using Randomized Axelerated Maximum Likelihood (RAxML) v8.2.12 (Stamatakis 2014). RAxML inferred trees in sliding windows (100kb of length, 20kb of slide) using a generalized time-reversible (GTR) CAT model with Lewis ascertainment bias correction. Twisst then estimated the weighting of each taxon topology (defined as the fraction of all unique population sub-trees) per window.

Finally, we calculated population specific *F*_ST_ from allele sharing matrices (β; (Weir and Goudet 2017; Goudet et al. 2018) in sliding windows of 100kbp with 20kbp steps, using hierfstat. For each matrix, we calculated i) the mean allele sharing for pairs of individuals from islands and ii) the mean allele sharing for pairs of individuals, one from an island and the other from the mainland, from which we obtain the islands specific *F*_ST_. We shall refer to this estimate as *F*_ST_ island-specific (*F*_ST_^Can^; see Sup. Fig. 1 for a schematic representation). This method allowed us to identify genomic regions of high differentiation exclusively on the islands with no confounding effect from the mainland (see also Weir et al. 2005).

From the genome wide scans, we identified peaks of differentiation with at least two overlapping windows of *F*_ST_^Can^ higher than 5 standard deviations (SD) from the mean (threshold: 0.377), and extracted the genes in these regions from the NCBI’s annotation of the reference genome (RefSeq accession: GCF_018691265.1). The gene list was then fed to ShinyGO v0.61 (Ge et al. 2020) to investigate potential enrichment of molecular pathways.

#### Eastern vs Western Canary Islands

On a second stage, to investigate potential genomic signals of differentiation, putatively linked with ecological adaptation to their distinct niches, we contrasted the two insular lineages (EC against WC). We estimated pairwise *F*_ST_, *d*_xy_ and π as for the island-mainland comparison described above.

Then, as for the island-mainland comparison above, we used hierfstat to calculate population specific *F*_ST_ in sliding windows of 100kbp with 20kbp steps based on a dataset including only the insular individuals. For each island, we consequently identified genomic regions with at least two overlapping windows of pairwise *F*_ST_ higher than 5 SD from the mean (regions highly differentiated between islands, threshold: 0.157) and above the 99^th^ quantile of each population’s F (regions highly similar within each island, respective threshold for *F*_ST_^EC^= 0.404 and for *F*_ST_^WC^ =0.393) and extracted the genes in these regions as described above. This yielded two genes lists, one per island, which were input to ShinyGO as above.

### Climatic niche analysis

To assess whether the barn owl populations of eastern and western Canaries occupy different climatic niches, we used the Outlying Mean Index (OMI) approach (Dolédec et al. 2000) as implemented in the R package ade4 v.1.7-16. Observation points for barn owls were compiled from three different sources: Global Biodiversity Information Facility (GBIF), samples sequenced in this study and in Burri et al. (2016). We kept records only from the islands sampled in this study, namely Tenerife (*T. alba*; N=79) and Lanzarote and Fuerteventura (*T. a. gracilirostris*; N=34 and N=39, respectively).

Climatic variables for the Canary Islands were extracted from the WorldClim database (Hijmans et al. 2005) at 30 sec resolution (approximately 1 km^2^) using the R package rbioclim (Exposito-Alonso 2017). Redundant variables (correlation of 1) were trimmed and the final model was run with the following 13 variables: Mean Diurnal Range (BIO2), Isothermality (BIO3), Temperature Seasonality (BIO4), Max Temperature of Warmest Month (BIO5), Min Temperature of Coldest Month (BIO6), Temperature Annual Range (BIO7), Mean Temperature of Driest Quarter (BIO9), Annual Precipitation (BIO12), Precipitation of Driest Month (BIO14), Precipitation Seasonality (BIO15), Precipitation of Warmest Quarter (BIO18), Precipitation of Coldest Quarter (BIO19).

## Results

### Population structure and genetic diversity

The overall *F*_ST_ in our dataset was 0.0698. Individual ancestry analyses with sNMF distinguished four genetic clusters, separating each population into its own cluster, except Morocco (MA) and Portugal (PT) that clustered together (Fig. 1a). Similarly, PCA clustering clearly grouped individuals according to their population, except PT and MA (Fig. 1c). The first axis opposed the insular populations to the mainland, as did sNMF K=2 (Sup. Fig. 2), with Eastern Canary (EC) and Israel (IS) at each extreme. The second axis contrasted the two insular populations EC and Western Canary (WC), as in K=3 (Sup. Fig. 2), and finally the third axis segregated the mainland groups with PT and MA opposed to IS, as in K=4 (Fig. 1a, 1c). Three individuals from EC (one from Fuerteventura and two from Lanzarote) showed small ancestry levels from both WC and PT in sNMF and were placed intermediately on axes 1 and 2 of the PCA. In the PCA including the European individuals, the first axis remained qualitatively the same with EC and WC together in opposition to IS (Sup. Fig. 3). Axes 2 and 3 switched positions, with MA, PT and the rest of Europe being first isolated from the rest and only then the two Canary split from each other (Sup. Fig. 3). In terms of differentiation, pairwise *F*_ST_ were highest between both islands and IS (EC-IS 0.092; WC-IS 0.088) as well as between islands (WC-EC 0.084). The mainland populations were overall less differentiated (all below 0.04), with PT and MA having the lowest *F*_ST_ with all other populations in accordance with their central position on the PCA (all below 0.06 for PT and 0.061 for MA), and the lowest between themselves (0.007) (Table 1).

**Table 1.** Genetic diversity and population differentiation of barn owls from the Mediterranean Basin and the Canary Islands. Right-hand-side of the table shows the matrix of population pairwise *F*_ST_.

| Population | Abbrev. | N | #PA | #rPA | $H_o$ | $F_{IS}$ | $F_{IT}$ | $F_{ST}$ | EC | WC | MA | PT | IS |
| --- | --- | --- | --- | --- | --- | --- | --- | --- | --- | --- | --- | --- | --- |
| Eastern Canary | EC | 10 | 324458 | 265175 (3829) | 0.161 (0.013) | -0.015 (0.08) | 0.10 (0.07) | EC | 0.112 | 0.081 | 0.061 | 0.060 | 0.088 |
| Western Canary | WC | 9 | 288313 | 288313 | 0.167 (0.006) | -0.030 (0.04) | 0.07 (0.03) | WC | 0.081 | 0.105 | 0.055 | 0.054 | 0.083 |
| Morocco | MA | 3 | 234524 | 692508 (13287) | 0.190 (0.001) | -0.04 (0.01) | -0.06 (0.01) | MA | 0.061 | 0.055 | -0.013 | 0.007 | 0.034 |
| Portugal | PT | 9 | 669889 |  | 0.182 (0.008) | -0.013 (0.04) | -0.01 (0.04) | PT | 0.060 | 0.054 | 0.007 | 0.014 | 0.039 |
| Israel | IS | 9 | 703392 | 703392 | 0.180 (0.002) | -0.036 (0.04) | -0.004 (0.01) | IS | 0.088 | 0.083 | 0.034 | 0.039 | 0.050 |

N – number of sampled individuals; #PA – private alleles in each population; #rPA – private alleles (SD) rarefied to 9 individuals per population (note that PT and MA were merged); $H_o$ – mean observed heterozygosity (SD); $F_{IS}$ – population level inbreeding coefficient (SD); $F_{IT}$ : mean individual inbreeding coefficient relative to the meta-population (SD). $F_{ST}$ : pairwise and population specific $F_{ST}$ .

Treemix yielded a population tree highly resembling the first PCA axis, with two main branches splitting from the IS root, one towards PT and MA and the second to WC and EC (Sup. Fig. 4). Most of the variance was explained by the tree with no migration (0.993). Nonetheless, the tree with two migration events displayed the highest likelihood (162.284), with one migration edge from the base of WC towards the tip of PT (edge weight 0.33) and the other from the base of PT towards the tip of EC (edge weight 0.13). Adding more than two migration events to the tree did not improve its fit to the data.

Overall, mainland populations presented higher genetic diversity than the insular ones (Table 1). Nonetheless, both insular populations showed over 280’000 private polymorphic sites (Table 1). Accordingly, individual relatedness was higher within and between insular populations (Sup. Fig. 5), and PT had the lowest within population relatedness. All populations showed signs of random mating with *F*_IS_ close to zero but slightly negative as expected of dioecious species (Balloux 2004), while the inbreeding levels of insular barn owls relative to the whole set of populations (*F*_IT_) were higher than those on the mainland (Table 1), a reflection of their isolation from the continent.

### Island vs mainland genomic comparisons

In order to measure the differentiation between islands and mainland individuals, we computed two distinct *F*_ST_ along the genome (see methods for details). *F*_ST_^Can^ allowed the detection of regions of high similarity in the islands as a whole, whereas *F*_ST_^ILvsML^ produced less clear results with population-specific allelic frequencies driving the overall signal instead of the defined groups. For example, *F*_ST_^ILvsML^ yielded a peak of differentiation between the islands and the mainland which, upon closer inspection of pairwise comparisons of the populations, turned out to be due to differences between EC and the mainland only, whereas WC did not produce the same signal (Sup. Fig. 6). The genomic comparisons of diversity and divergence between insular and mainland barn owls based on the *F*_ST_^Can^ yielded 48 regions (478 100kb overlapping windows) of high differentiation including 100 genes (Fig. 2a). ShinyGO analyses identified an enrichment of five functional categories related to morphogenesis in humans (Sup. Table 2). The largest enriched pathway – anatomical structure morphogenesis – included 25 of the genes in regions of high genomic differentiation. The remaining three categories – anatomical structure formation involved in morphogenesis, tube morphogenesis and blood vessel morphogenesis – were subsets of the largest pathway, including 15, 14, 12 and 10 of the 25 genes, respectively.

**Figure 2.**
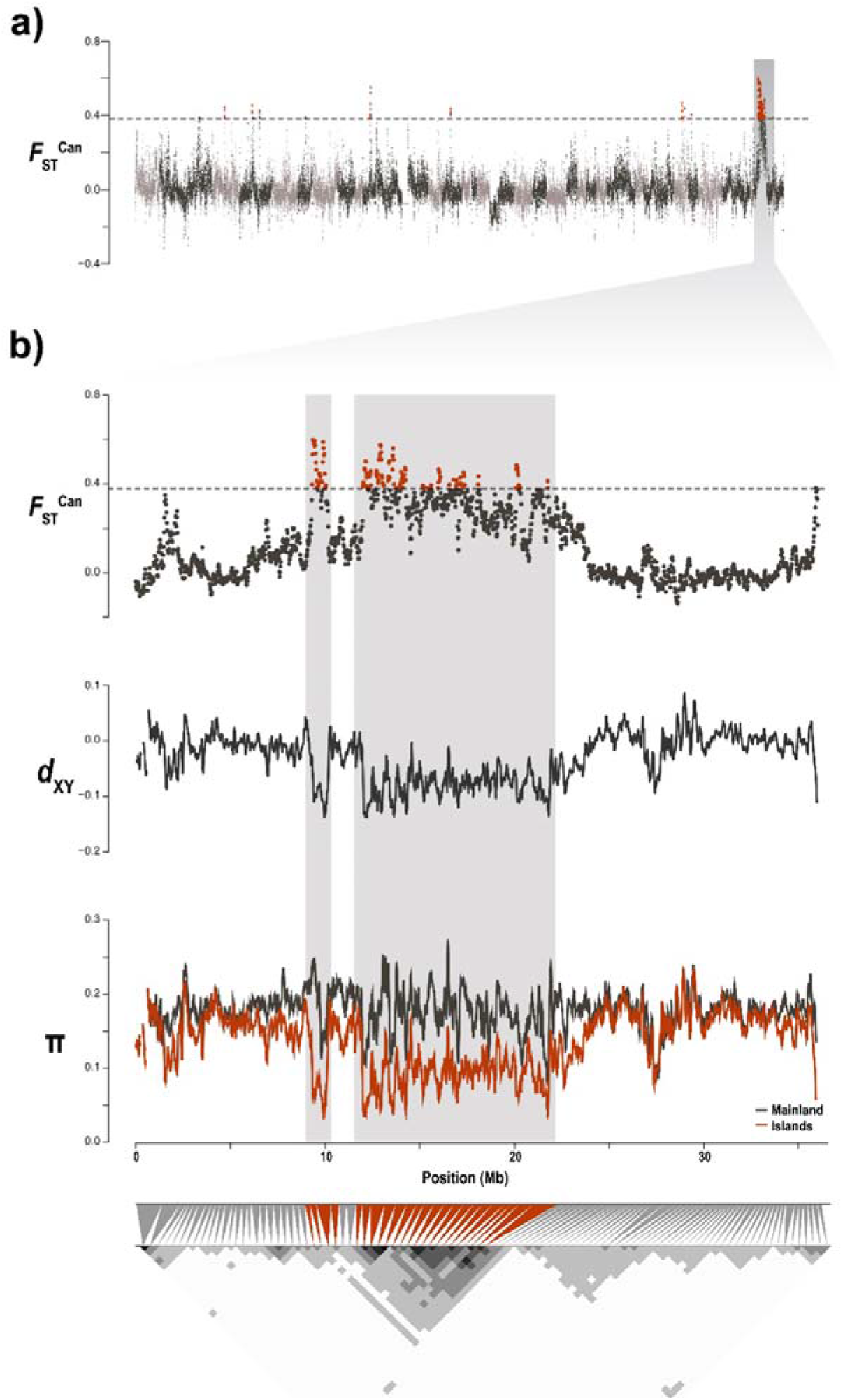
Genomic landscape of differentiation between insular and mainland barn owls. **a)** Genome-wide beta comparison between individuals from the Canary Islands (EC & WC) and from the mainland (PT & IS). Each dot represents a 100kb window. Dashed line indicates the 5 SD threshold used to identify genomic regions of high differentiation, emphasised in red. Alternating grey colours denote different scaffolds. Shaded vertical bar highlights Scaffold 100006. **b)** Zoom on Scaffold 100006 and, in particular, the ~15 Mb long highly differentiated genomic region (background shading) over windows of 100 kb. From top to bottom, we see in this region a high *F*_ST_^Can^ between insular and mainland barn owls, low absolute distance (*d*_XY_) between both islands (scaled with mean=0) and reduction of nucleotide diversity (π) among insular individuals (red line) compared to the mainland (black line). The bottom triangular matrix shows pairwise LD (*r*^2^) between groups of 100 SNP along the chromosome in insular owls. Darker pixels show higher LD. Grey triangles match each pixel in the matrix diagonal to the region it spans on the chromosome above. Red triangles indicate pixels that overlap the region of high differentiation.

The largest of the peaks encompassed 15Mb of Super-Scaffold_1000006. Of all the genes found in peaks of differentiation, 87 out of 100 were within this region. Here, insular owls showed a strong decrease in relative diversity compared to the mainland (*F*_ST_^Can^), as well as a drop of absolute divergence (*d*_xy_) between each other and nucleotide diversity (π). The region showed strong LD (*r*^2^) in all insular owls between neighbouring variants compared to the rest of the scaffold (Fig. 2b), which was not the case among continental ones (Sup. Fig. 7). In addition, *F*_ST_ was higher between island and mainland owls (Sup Fig. 6). Twisst showed roughly similar proportions of each tree along the genome, except for this region where there was a higher than average proportion of trees that joined EC and WC (Sup. Fig. 8). Of the 25 genes of the morphogenesis pathway, 18 were found in this closely-linked region of Super_Scaffold_1000006 (21% of the total 87 genes of the region).

### Eastern vs Western Canary Islands

#### Climatic niche analysis

To investigate the genomic and ecological differentiation between island populations (EC and WC), we first depict different climate in the two set of islands based on barn owl observations in the Canary Islands. The climatic niche analyses (OMI) yielded two axes that explained the climatic variability in our study area (Fig. 3b), with the first axis (OMI1) explaining nearly all of it (99.3%). OMI1 was positively correlated with temperature and negatively correlated with precipitation (Sup. Fig. 10). The eastern population *T. a. gracilirostris* occupied a narrow niche of high temperature, low precipitation and low seasonal and daily variability. *Tyto alba* in Tenerife occupied a broader niche that covered most of OMI1, including some of the niche of *T. a. gracilirostris*. OMI2 explained little of the variability (0.7%), spreading slightly each population’s niche without segregating them.

**Figure 3.**
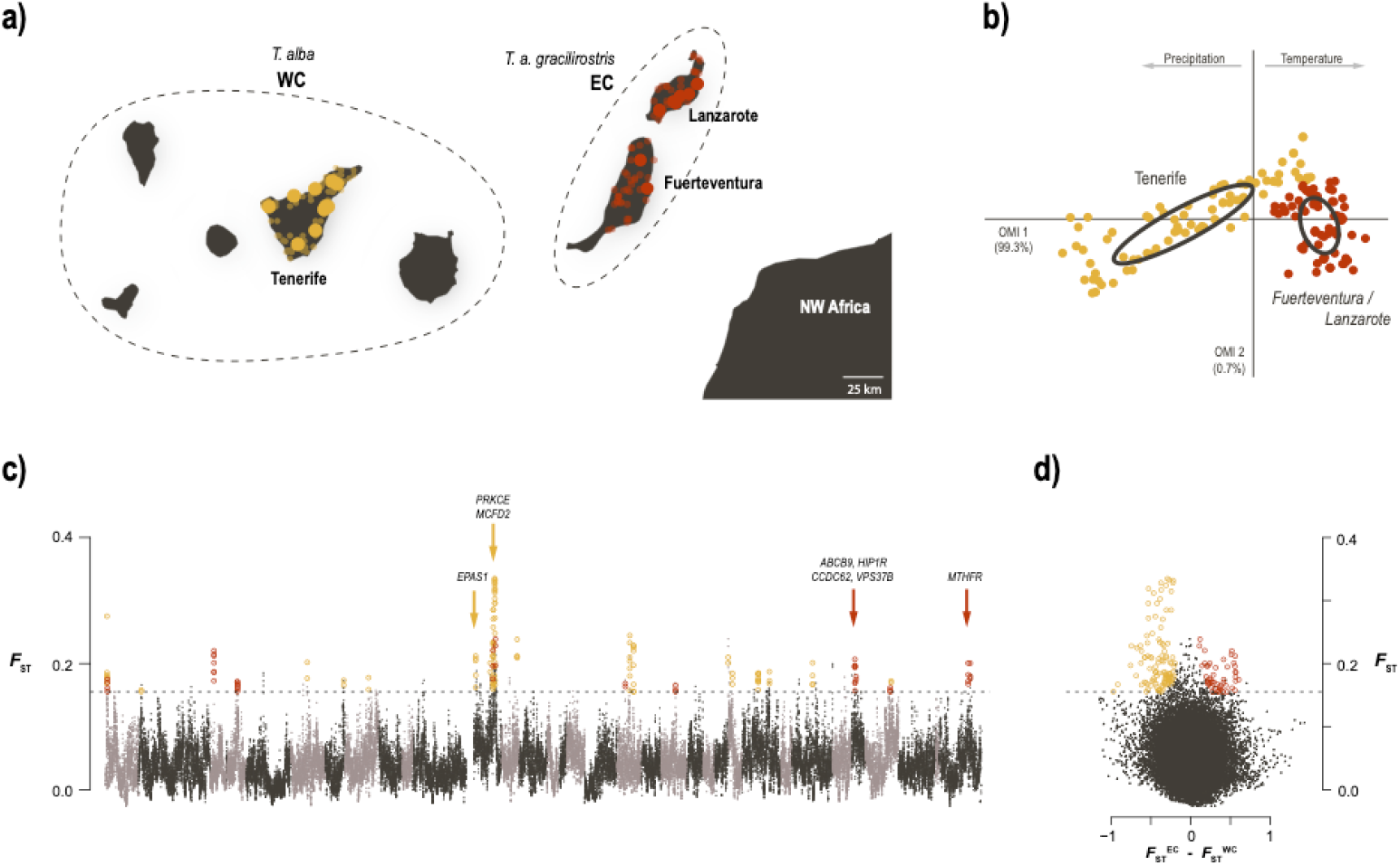
Ecological divergence in barn owls from the Canary Islands. **a)** Sampling location of Western (WC; yellow) and Eastern (EC; red) Canary individuals, off the coast of north-western Africa. Sampled islands are named. Large dots indicate individuals sampled for WGS and population genomics analyses; small transparent dots are barn owl observations used in niche analysis. Dashed lines group islands according to barn owl taxonomy. **b)** Climatic niche occupied by the two Canary populations. The first axis varies from cold and wet to hot arid environments. **c)** Genome-wide *F*_ST_ comparison between individuals from each Canary population. Each dot represents a 100kb window. Alternating grey colours denote different chromosomes. Dashed line indicates the 5 SD *F*_ST_ threshold used to identify genomic regions of high differentiation overlaid with *F*_ST_^WC^, highlighted in yellow and *F*_ST_^EC^ in red. Arrows denote the location of genes linked with response to hypoxia in WC; and the cluster of 4 genes linked to morphological ratios and the main gene related to blood pressure in EC. **d)** Same *F*_ST_ windows and threshold as in c plotted against the difference between *F*_ST_^WC^ and *F*_ST_^EC^, clearly highlight the two sets of windows putatively under local adaptation.

#### Detection of differentially selected genomic regions

We then searched for regions under selection in each island, yielding to a total of 33 putatively adaptive regions (14 in EC and 19 in WC), with high differentiation (*F*_ST_) between island and increased relatedness within each island (*F*_ST_^EC^ and *F*_ST_^WC^), from which we obtained two lists of genes (Fig. 3c, 3d). For EC, there were 30 such genes (Sup. Table 3) while for WC there were 25 genes (Sup. Table 4). Enrichment analyses found no link to a specific GO pathway for either island population. Nonetheless, in each island there were clusters of genes with similar identified functions, namely body proportions and blood pressure in EC, and cellular response to hypoxia in WC.

## Discussion

Islands offer unique conditions for organisms to adapt and expand their niches. In the Canaries, an archipelago rich in endemics, the barn owl is one of the few raptors present and is thought to have diverged into a subspecies on the easternmost islands. Taking advantage of whole-genome sequences, we first investigate the origin of the population of barn owls in the Canary Islands, revealing that the latter are long-standing populations and allowing us to address their taxonomic classification. Using a new, more sensitive method based on excess of allele sharing within the islands, we detect 58 putatively locally adapted genomic segments, many of them grouped in a haplotype-like region seemingly under positive selection in the Canary Islands. A quarter of the tightly linked genes in this genomic region are enriched for genes contributing to a pathway of anatomical morphogenesis suggesting morphological adaptations to insularity. We also identify scattered genomic regions putatively locally adapted to either the eastern and western islands. For eastern Canaries, the identified genes in regions under positive selection belong to pathways linked to body proportions and blood pressure, consistent with the smaller owl size of this population living in a hot arid climate. In the western Canaries, barn owls from Tenerife display a potential signal of selection of genes related to hypoxia, a potential response to inhabiting an island with a steep elevation gradient.

### Barn owls from the Canary Islands and Mediterranean Basin

Our work shows that, while each population in the study area has its own unique genetic composition (Fig. 1), barn owls from the Canary Islands are distinct from those on the mainland surrounding the Mediterranean Basin. Indeed, clustering methods constantly opposed insular individuals to mainland ones (Fig. 1c; Sup. Fig. 2) and were supported by the drift-based trees of Treemix. Thus, overall, our current results indicate that the two insular lineages – Eastern and Western Canaries – have a common origin.

In terms of genetic diversity, the islands were less diverse than the mainland as expected and showed slightly higher levels of inbreeding relative to the whole set (Frankham 1997). However, they presented nearly 10-fold higher levels of private alleles than those reported for other islands in the Atlantic (British Isles; Machado et al. 2021) and in the Mediterranean Sea (Cyprus and Greek islands; Machado et al. 2021). Notably, the two Canaries insular populations, about 165 km apart, were more distant genetically from each other than either was from Portugal, over 1000 km away. Furthermore, they were more distant from each other even than WC was from any population in continental Europe in a recent genomic study (Cumer et al. 2021), as well as in an earlier one with microsatellites (Burri et al. 2016). The high private diversity could be partially explained by the small number of samples in north-western Africa, which may be masking any shared polymorphisms between the islands and the nearest mainland as private to the former. However, as we did sample PT and IS, the most diverse populations in the Western Palearctic assumed to meet in northern Africa (Cumer et al. 2021), much of this insular diversity is likely indeed private. Therefore, the Canaries appear to have been colonized much earlier than the other studied insular populations and have thus had the time for *in situ* mutations to accumulate in spite of genetic drift, which is supported by their higher divergence as well. These populations may also reflect a more ancient lineage from northern Africa that would have been replaced or simply evolved in a different direction and only the addition of samples from African mainland (on top of the three MA samples) would allow to fully depict the neutral history of these populations.

### Insularity

We found multiple evidence of adaptation common to both barn owl lineages of the Canary Islands. To do so, we used an estimator of population specific *F*_ST_ (Weir and Goudet 2017; Goudet et al. 2018) to identify genomic regions with an excess of shared ancestry in all insular individuals relatively to the mean shared ancestry between islands and mainland individuals. Being a moment estimator, it can process efficiently a large amount of genomic loci, compared to maximum likelihood or Bayesian estimators (e.g. Foll and Gaggiotti 2008). More importantly, it does not rely on the F-model, which assumes independent populations. This island specific *F*_ST_ (*F*_ST_^Can^) identified regions of increased relatedness between insular individuals compared to the averaged relatedness along the genome (i.e. regions in which all insular individuals resemble each other more than expected). This population specific, moment based and model-free estimator of *F*_ST_ provides a clearer result than classical pairwise *F*_ST_ scan, as it focuses on the diversity in the target population rather than taking an average over the set of populations, and should be a useful addition to the population genomic toolbox to detect nested signals of local adaptation, especially when there is substructure in the groups one wishes to compare. We thus considered regions highly similar in insular individuals as putatively under selection on the islands. The genomic landscape of *F*_ST_^Can^ differentiation between insular and mainland owls yielded 58 windows putatively under selection (Fig. 2a). Among these, a particularly large and clear peak of differentiation stood out. This region approximately 15 Mb in length was highly similar among insular individuals (Sup. Fig. 9) as shown by the accompanying drop in *d*_XY_ and π (Fig. 2b). Furthermore, the increased linkage between alleles in this region suggests that it is transmitted in a haplotype-fashion. Crucially, the fact that we do not see the slightest surge of LD in mainland individuals confirms that it is not a by-product of a region of low-recombination in this species (Sup. Fig. 7). Overall, we provide strong evidence of positive selection in this genomic region in Canary Islands owls, suggesting an adaptation to insularity (Fig. 2).

A fifth of the genes in this haplotype (18 out of 87), in conjunction with 7 other genes in potentially adaptative regions, significantly enriched the anatomical structure morphogenesis pathway (Sup. Table 2), a biological process related to the organisation and generation of anatomical structure during development. This suggests positive selection on some morphological trait on insular individuals. Given that there is evidence of gene flow from the mainland into both islands (see admixed individuals in Fig. 1, Sup Fig, S2), we propose two hypotheses to explain how selection might act on this haplotype. First, it could confer a significant advantage to individuals carrying it on the island and prevent those that do not carry it from reproducing or surviving. In this scenario, immigrants from the mainland not carrying this haplotype would reproduce less in both islands. In the second scenario, selection would happen on the migrants themselves before reaching the island if, for example, the haplotype facilitates long flight over large spans of water. It is widely accepted that, given how dispersal capacity is highly variable across species, even among birds, some are more prone to colonizing islands than others. Therefore, it is also conceivable that, within a species or population, some individuals are morphologically more predisposed or have better dispersal abilities than others. Since the barn owl generally avoids flying over open water, as demonstrated by its consistently higher differentiation on islands (Cumer et al. 2021; Machado et al. 2021, n.d.), this seems plausible. The absence of phenotypic measurements from the sequenced birds prevents us from establishing a link between phenotypes and genotypes on this data set, and we hence remain cautious on the speculation regarding the functional implications of this haplotype. Future work should verify the frequencies of this haplotype in a larger cohort as we only had 19 insular individuals in this study and, if possible, include detailed morphometric measurements to allow a GWAS-like approach.

### Ecological divergence

In the Canary archipelago, both the eastern islands and Tenerife have many specific endemic species across multiple taxa. This is generally attributed to their intrinsic characteristics driving ecological speciation, namely the arid and windy conditions of Lanzarote and Fuerteventura, and the elevation gradient of the Teide volcano (up to 3’718m a.s.l.) in Tenerife. We quantified the climatic differences between the two environments with a niche analysis based on reported barn owl observations, and show that indeed barn owls occupy significantly different niches on each group of islands (Fig. 3b). In the east, they are found on unvaryingly hot and dry locations, whereas Tenerife covers a wide range of temperature and precipitation (Fig. 3b).

From the genomic data, we found evidence of local adaptation on both insular populations (Fig. 3c, 3d). The eastern population had more genomic regions, and more genes, potentially under selection compared to the west (30 and 25 genes, respectively). Although no significant pathway enrichment was detected in either population, there were clear groups of genes with similar known functions in the putatively adapted regions. In the eastern population there were two such groups. The first, composed of 7 genes, has significant links to body size and proportions in humans (see Sup. Table 3 and references therein). Among these, a specific set of 4 genes – *HIP1R, CCDC62, VPS37B and ABCB9* – is tightly clustered in the barn owl genome (Fig. 3c). These four genes have been linked to body height, body-mass-index and other body measurement ratios (Turcot et al. 2018; Kichaev et al. 2019; Vuckovic et al. 2020; Zhu et al. 2020), and may thus play a role in the smaller size of barn owls in the eastern population, which would then be adaptive. The second group of genes in regions potentially under selection, includes 11 genes related to numerous blood parameters (Sup. Table 3), a similar signal to that seen in chickens adapted to hot arid environments (Gu et al. 2020). In particular, the gene *MTHFR* has extensive connections to blood pressure (Newton-Cheh et al. 2009; Ehret et al. 2016; Liu et al. 2016; Surendran et al. 2016; Hoffmann et al. 2017; Wain et al. 2017; Kulminski et al. 2018; German et al. 2020), a trait known to vary with environmental temperature in mammals (Halonen et al. 2011) and birds (Darre and Harrison 1987), potentially suggesting barn owls have circulatory systems adapted to the hot and dry conditions of the eastern Canaries.

In the western island population, regions putatively under selection contained an interesting group of 4 genes - *EPAS1, PRKCE, MCFD2* and *FZD8* - with links to red blood cells, haemoglobin density and cellular response to hypoxia (i.e. low levels of oxygen; Sup. Table 4; Astle et al. 2016; Kichaev et al. 2019; Chen et al. 2020; Oskarsson et al. 2020; Vuckovic et al. 2020). Red blood cells, and the haemoglobin within, are responsible for transporting oxygen in the body and are direct targets of selection at high elevation (O’Brien et al. 2020). The gene *EPAS1* in particular, is well known for being involved in adaptation to high altitude environments across vertebrates (Witt and Huerta-Sánchez 2019). These genes hints at a possible adaptation of barn owls to higher altitude in Tenerife. On this small island, barn owls are mostly concentrated at lower altitude (i.e. along the coast – 0 to 300m a.s.l.) and their repartition is limited in the central part of the island due to the presence of the high mountains at about 2000m a.s.l., that culminates with the colossal peak of Teide at 3715m a.s.l. (Siverio 1998). This is consistent with the known limited ability of this bird to live at high altitudes (Machado et al. 2018; Cumer et al. 2021). However, the observation of individuals up to 1200m a.s.l. and the presence of breeding couples at moderate altitudes (600 to 1200m a.s.l., Siverio 1998), support the hypothesis of an ongoing adaptation to higher altitude in this insular population. This hypothesis, which deserves further investigations, is also consistent with observations made in other populations in warm climates; where local barn owl populations expand their range by adapting to slightly higher altitudes (Romano et al. 2020).

Considering its wide distribution, even accounting for phenotypic plasticity, barn owls’ capacity to adapt to a variety of prey and environments is unquestionable. As such, detecting signals of local adaption in the Canary Islands is not surprising. Indeed, with islands generally being species-poor, the species that do inhabit them have to adapt to different or broader niches via ecological divergence (Losos and Ricklefs 2009). This is especially true of volcanic islands that arise isolated and uninhabited, in contrast to those intermittently connected to the mainland and more easily colonized. Moreover, the community of birds of prey in the Canary archipelago includes less than half the species found in the nearby mainland (8 diurnal and 2 nocturnal species; Martín and Lorenzo, 2001; Rodríguez et al. 2018), likely due to the lack of suitable habitat, preys and/or the limited surface. A by-product of this is the reduction of inter-specific competition, which could have allowed the barn owl to maintain population sizes just large enough on the islands through time for selection to act and potentially expand its niche to better exploit the insular environment. Our results suggest this is happening in parallel on each island (Fig. 3c, 3d), consistent with their different niches (Fig. 3b) and relative genetic isolation, producing two distinct ecomorphs, adapted to distinct ecosystems.

### Insular subspecies

The eastern islands of Lanzarote and Fuerteventura, and adjacent islets (Fig. 3a) are home to the barn owl subspecies *T. a. gracilirostris*. This classification is based on its smaller size and even on colouration pattern, although the latter is contested by ornithologists and inconsistent with reported phenotypical measurements (Burri et al. 2016). The reduction in size is actually a common pattern in insular barn owls (Romano et al. 2021), and could be an adaptation to nesting in very small cavities (i.e. cracks in lava walls) and/or to better navigate the strong winds in the eastern islands (Siverio 2007). The genomic data presented here is consistent with this population forming an endemic subspecies. It has diverged considerably from the mainland, with higher differentiation levels than barn owls from any other studied island in the Western Palearctic (Machado et al. 2021, n.d.). Moreover, we show it carries high levels of private genetic diversity and multiple genomic regions showing signs of local adaptation (Fig. 3c).

In contrast, barn owls from Tenerife and the remaining islands are considered to belong to the nominal *T. alba* found also on the mainland surrounding the Mediterranean Basin (Fig. 3a). However, it too has considerably diverged from the mainland and shows signs of potential ongoing adaptation in Tenerife and its elevation gradient (see previous section). Furthermore, it clusters with the other insular Canary population rather than the mainland (Fig. 1). While it is not the aim of this study to evaluate what constitutes a subspecies, we provide evidence that the Tenerife population is diverging significantly from its founding population, both neutrally and adaptively, albeit at a slower pace than the eastern population.

The reasons why the eastern population is more divergent than the western, a puzzling fact considering it is closer to the mainland, are not yet fully resolved. Neutral divergence in *F*_ST_ between these two insular populations suggest they are the result of two independent colonisation events rather than a strict east-to-west progression as described for other taxa (Juan et al. 2000). Although the islands themselves emerged from east to west, Tenerife is at least 11 million years old, twice the inferred time of formation of the *T. alba* species (Uva et al. 2018) and thus available for colonisation at the time. Nonetheless, an earlier settlement of the eastern islands would have given more time for both genetic drift and selection to promote divergence. Alternatively, a very small population size in the east, consistent with current census data (Palacios 2004; Siverio 2007), could account for the stronger drift in a scenario of simultaneous colonization. However, it would strongly hinder local adaptation, making it a less likely hypothesis since we identified more regions putatively under selection in this population. Further work, including an extensive sampling of all island populations, as well as demographic modelling, would be needed to resolve this intriguing pattern and refine the history of the barn owl in the Canary archipelago.

## Conclusion

Due to their intrinsic characteristics, islands house numerous endemics making them ideal systems to study the bases of ecological divergence. We provide empirical evidence that both neutral and adaptive evolutionary mechanisms shaped divergence from the mainland in barn owls from the Canary archipelago. Our results show clear signs of genome-wide differentiation (i.e. neutral), a combination of mutations (high private diversity; Table 1) and drift (high *F*_ST_ and *F*_IT_), consistent with theoretical expectations for populations established and isolated long ago despite some admixture. We also identify signals of local adaptation to common insular conditions (Fig. 2), as well as to each island’s niche, creating ecomorphs (Fig. 3). While the history and functional effect of the putatively adapted genomic regions identified here deserve further investigation, these observations highlight how selection can still act on small isolated populations. This study illustrates the capacity of a widespread bird to adapt to the local ecological conditions of small islands, an adaptative capacity which may prove essential in facing a changing global climate.

## Supporting information

Supplementary Informations

## Acknowledgements

We are grateful to the following institutions and individuals that provided samples or aided in sampling to our study: Karim Rousselon, Gustavo Tejera, Sylvain Antoniazza, Reto Burri, Aurelio Martín, David P. Padilla, the “Association Marocaine pour la Protection des Rapaces AMPR”, the Natural History Museum of Tenerife and the Wildlife Rehabilitation Centre La Tahonilla (Cabildo Insular de Tenerife). We thank Céline Simon and Guillaume Dumont for their valuable assistance with molecular work. The Gobierno de Canarias provided all the corresponding permits for the collection and inter-island transport of samples. This study was funded by the Swiss National Science Foundation with grants 31003A-138180 & 31003A_179358 to JG and 31003A_173178 to AR.

## Data Accessibility

The raw Illumina reads for the whole-genome sequenced individuals are available in BioProjects PRJNA700797, PRJNA727977 and PRJNA774943. Sampling locations used for niche modelling are given in appendix 1.

## Author Contributions

APM, TC, AR, JG designed this study; FS, MC, IR, RL provided samples; APM produced whole-genome resequencing libraries; TC and APM conducted the analyses; APM and TC led the writing of the manuscript with input from all authors.

