## Supplementary Informations for "Genomic bases of insularity and ecological divergence in barn owls (*Tyto alba*) of the Canary Islands"

### Supplementary Tables

**Sup. Table 1** – Description of samples used in this study.

| # | Pop | ID | Location | Year | Tissue | Sex | Ref |
|---|---|---|---|---|---|---|---|
| 1 | East Canary | EC01 | Lanzarote | 2010 | muscle | Female | 1 |
| 2 | East Canary | EC02 | Lanzarote | 2010 | muscle | Female | 1 |
| 3 | East Canary | EC03 | Lanzarote | 2012 | muscle | Female | 1 |
| 4 | East Canary | EC04 | Lanzarote | 2012 | muscle | Female | 1 |
| 5 | East Canary | EC05 | Lanzarote | 2012 | muscle | Male | 1 |
| 6 | East Canary | EC06 | Lanzarote | 2012 | muscle | Male | 1 |
| 7 | East Canary | EC07 | Lanzarote | 2007 | feather | Male | 1 |
| 8 | East Canary | EC08 | Lanzarote | 2008 | feather | Female | 1 |
| 9 | East Canary | EC09 | Fuerteventura | 2007 | feather | Male | 1 |
| 10 | East Canary | EC10 | Fuerteventura | 2007 | feather | Female | 1 |
| 11 | West Canary | WC01 | Tenerife | 1905 | muscle | Male | 2 |
| 12 | West Canary | WC02 | Tenerife | 2003 | muscle | Female | 2 |
| 13 | West Canary | WC03 | Tenerife | 2003 | muscle | Female | 2 |
| 14 | West Canary | WC04 | Tenerife | 2003 | muscle | Male | 2 |
| 15 | West Canary | WC05 | Tenerife | 2005 | muscle | Male | 2 |
| 16 | West Canary | WC06 | Tenerife | 2005 | muscle | Female | 2 |
| 17 | West Canary | WC07 | Tenerife | 2006 | muscle | Male | 2 |
| 18 | West Canary | WC08 | Tenerife | 2006 | muscle | Male | 2 |
| 19 | West Canary | WC09 | Tenerife | 2006 | muscle | Male | 2 |
| 20 | Portugal | PT01 | Pombal | 2013 | feather | Female | 3 |
| 21 | Portugal | PT02 | Coruche | 2013 | feather | Male | 3 |
| 22 | Portugal | PT03 | Évora | 2013 | feather | Male | 3 |
| 23 | Portugal | PT04 | Coruche | 2012 | feather | Female | 3 |
| 24 | Portugal | PT05 | Nazaré | 2013 | feather | Female | 3 |
| 25 | Portugal | PT06 | Porto de Moós | 2013 | feather | Female | 3 |
| 26 | Portugal | PT07 | Setúbal | 2012 | feather | Female | 3 |
| 27 | Portugal | PT08 | Fátima | 2013 | feather | Female | 3 |
| 28 | Portugal | PT09 | Santarém | 2013 | feather | Male | 3 |
| 29 | Israel | IS01 | Lachish | 2005 | blood | Female | 2 |
| 30 | Israel | IS02 | Beit Shean | 2005 | blood | Male | 2 |
| 31 | Israel | IS03 | Hula | 2005 | blood | Female | 2 |
| 32 | Israel | IS04 | Beit Shean | 2005 | blood | Female | 2 |
| 33 | Israel | IS05 | Beit Shean | 2005 | blood | Male | 2 |
| 34 | Israel | IS06 | Beit Shean | 2005 | blood | Female | 2 |
| 35 | Israel | IS07 | Beit Shean | 2005 | blood | Female | 2 |
| 36 | Israel | IS08 | Hula | 2005 | blood | Female | 2 |
| 37 | Israel | IS09 | Hula | 2005 | blood | Female | 2 |
| 38 | Syngapore | SGP | Singapore | 2013 | soft tissue | Male | 3 |
| 39 | USA | USA | San Diego, California | 2015 | soft tissue | Female | 3 |

(*1*) GenBank BioProject PRJNA727977

(*2*) GenBank BioProject PRJNA774943

(*3*) GenBank BioProject PRJNA700797

**Sup. Table 2** – ShinyGO pathway enrichment results for genes in putatively adapted genomics regions in the insular Canarias populations. Only pathways enriched by more than three genes are displayed. All listed genes are located in the haplotype-like region in Figure 2, except those marked with †.

| Enrichment FDR | Genes in list | Total genes | Functional Category | Genes |
|---|---|---|---|---|
| 0.0443 | 15 | 1164 | Anatomical structure formation involved in morphogenesis | GNG5 ANXA2† F3 RORA† VAV3 CCN1 PRKACB DDAH1 WDR72† S1PR1 COL11A1 ADAM12† TGFBR3 GLMN DLC1† |
| 0.0443 | 25 | 2785 | Anatomical structure morphogenesis | PLPPR4 NEXN GNG5 ANXA2† TGFBR3 F3 CCN1 COL11A1 RORA† PALMD OLFM3 VAV3 BCAR3 PRKACB BARHL2 DDAH1 NTNG1 WDR72† S1PR1 ADAM12† FGD5† DLC1† DOCK1† GLMN USP33 |
| 0.0443 | 14 | 1077 | Circulatory system development | GNG5 ANXA2† TGFBR3 F3 COL11A1 ACADM VAV3 CCN1 DDAH1 S1PR1 GLMN RORA† ADAM12† DLC1† |
| 0.0460 | 12 | 860 | Tube morphogenesis | ANXA2† F3 VAV3 CCN1 PRKACB DDAH1 S1PR1 RORA† ADAM12† TGFBR3 GLMN DLC1† |
| 0.0486 | 10 | 603 | Blood vessel morphogenesis | ANXA2† F3 VAV3 CCN1 DDAH1 S1PR1 RORA† ADAM12† TGFBR3 GLMN |

**Sup. Table 3** – List of genes in putatively adapted genomic regions in the Eastern Canary population, grouped per location on the barn owl genome. Associated phenotypes are provided for each gene when available. Human-specific behavioural phenotypes are not reported (for example, alcohol consumption). Genes marked with † have phenotypes related to body size and proportions, and ‡ with blood parameters.

| Super-Scaffold | Gene | Phenotype | References |
|---|---|---|---|
| 3 | LRTM1† | Body height | (*4*, *5*) |
| 5 | LAS1L | - | |
| | MSN | Alopecia | (*6*) |
| | HEPH | Alopecia | (*6*) |
| | EDA2R | Alopecia | (*6*) |
| 16 | SRBD1 | - | |
| | DAAM2†‡ | Body height | (*4*, *7*) |
| | | Platelet count | (*8*) |
| 23 | MMAA‡ | Hemoglobin measurement | (*9*, *10*) |
| 44 | PITPNM2‡ | Platelet count | (*11*) |
| | | Reticulocyte count | (*10*) |
| | | BMI | (*4*) |
| | ARL6IP4 | - | |

| | | | |
|---|---|---|---|
| | OGFOD2 | - | |
| | ABCB9† | Waist-hip ratio | (*4*) |
| | | BMI-adjusted waist-hip ratio | (*12*) |
| | VPS37B† | BMI-adjusted waist-hip ratio | (*12*) |
| | HIP1R†‡ | BMI | (*13*) |
| | | BMI-adjusted waist-hip ratio | (*12*) |
| | | Waist-hip ratio | (*4*) |
| | | Body height | (*4*) |
| | | Platelet count | (*10*) |
| | CCDC62†‡ | BMI-adjusted waist-hip ratio | (*12*) |
| | | Platelet count | (*10*) |
| | MPHOSPH9‡ | Platelet count | (*9*, *10*, *14*) |
| | MTRFR | - | |
| | CDK2AP1 | - | |
| | SBNO1‡ | Lymphocyte count | (*10*) |
| 20000042 | P3H1 | - | |
| | CLDN19 | - | |
| | YBX1 | - | |
| | PPIH | - | |
| | CCDC30‡ | Systolic blood pressure | (*15*) |
| | | Pulse pressure | (*15*, *16*) |
| | PPCS | - | |
| | PLOD1‡ | Platelet count | (*14*) |
| | KIAA2013† | Waist-hip ratio | (*5*) |
| | CLCN6‡ | Diastolic blood pressure | (*17*, *18*) |
| | | Systolic blood pressure | (*4*, *17*, *18*) |
| | | Pulse pressure | (*18*) |
| | MTHFR‡ | Mean arterial pressure | (*19*) |
| | | Diastolic blood pressure | (*17*, *18*, *20*, *21*) |
| | | Systolic blood pressure | (*18*, *20–23*) |
| | | Pulse pressure | (*18*) |
| | | Platelet count | (*8*, *10*) |
| | | Blood pressure | (*20*) |
| | | Hypertension | (*24*) |
| | | Mean corpuscular volume | (*4*, *8*, *10*) |
| | | Erythrocyte count | (*4*, *10*) |

Sup. Table 4 – List of genes in putatively adapted genomic regions in the Western Canary population, grouped per location on the barn owl genome. Associated phenotypes are provided for each gene when available. Human-specific behavioural phenotypes are not reported (for example, alcohol consumption). Genes marked with † are linked to red blood cells and haemoglobin measurements, and with ‡ to other blood parameters.

| Super-Scaffold | Gene | Phenotype | References |
|---|---|---|---|
| 2 | WNT11 | Platelet count | (*9*, *10*) |
| 7 | GRB14 | Hemoglobin measurement | (*9*) |
| | | Erythrocyte count | (*10*) |
| | COBLL1 | BMI-adjusted waist-hip ratio | (*5*, *12*) |
| | | Erythrocyte count | (*14*) |
| | | Systolic blood pressure | (*4*, *14*, *16*) |
| 8 | OTUD7B | Body height | (*25*) |
| | | BMI-adjusted waist-hip ratio | (*12*) |
| | MTMR11 | Body height | (*4*, *5*) |
| | SF3B4 | - | |
| | SV2A | Body height | (*26*, *27*) |
| | BOLA1 | - | |
| | HJV | - | |
| | POLR3GL | - | |
| | ANKRD34A | Leucocyte count | (*9*) |
| | RBM8A | - | |
| | LIX1L | - | |
| 16 | SPTLC3 | - | |
| | TRERF1 | Leucocyte count | (*4*, *10*, *14*) |
| | UBR2 | - | |
| | CAMKMT | Body height | (*4*, *25*, *26*) |
| | TMEM247 | - | |
| | EPAS1† | Erythrocyte count | (*4*) |
| | | Hematocrit | (*10*) |
| | | Hemoglobin measurement | (*10*, *28*) |
| | | High altitude adaptation | (*29*, *30*) |
| | | PR interval | (*31*) |
| | PRKCE† | Hematocrit | (*8*, *10*, *14*, *32*–*36*) |
| | | Erythrocyte count | (*4*, *8*, *10*, *14*, *33*, *34*, *36*, *37*) |
| | | Hemoglobin measurement | (*8*, *10*, *14*, *32*–*36*, *38*) |
| | MCFD2† | Hematocrit | (*10*, *14*) |
| | | Erythrocyte count | (*10*, *14*) |
| | | Hemoglobin measurement | (*10*, *14*, *38*) |
| | | Body height | (*4*, *5*) |
| | LRFN2 | BMI | (*4*, *12*, *39*) |
| 38 | NXPH1 | - | |
| | GJD4‡ | PR interval | (*40*) |
| | FZD8† | Hematocrit | (*8*) |
| | | Erythrocyte count | (*8*, *10*) |
| | | Hemoglobin measurement | (*4*, *10*) |

#### Supplementary Figures

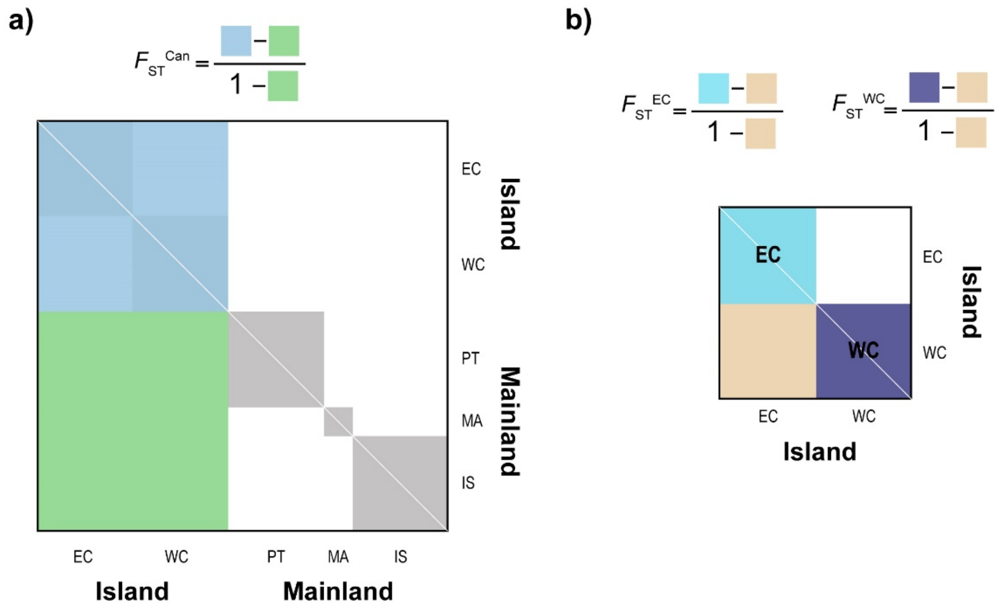

**Sup. Fig. 1** – Graphical representation of the calculation of *F*β from individual relatedness matrices to compare **a)** insular and mainland individuals and **b)** individuals from each island. Coloured squares in formulas on top, represent the mean of the same-colour section of the matrix underneath, calculated in windows of 100kb along the genome. Note that in **b)** the matrix is smaller as it does not include any owl from the mainland. In both matrices, the diagonal, in white, is empty.

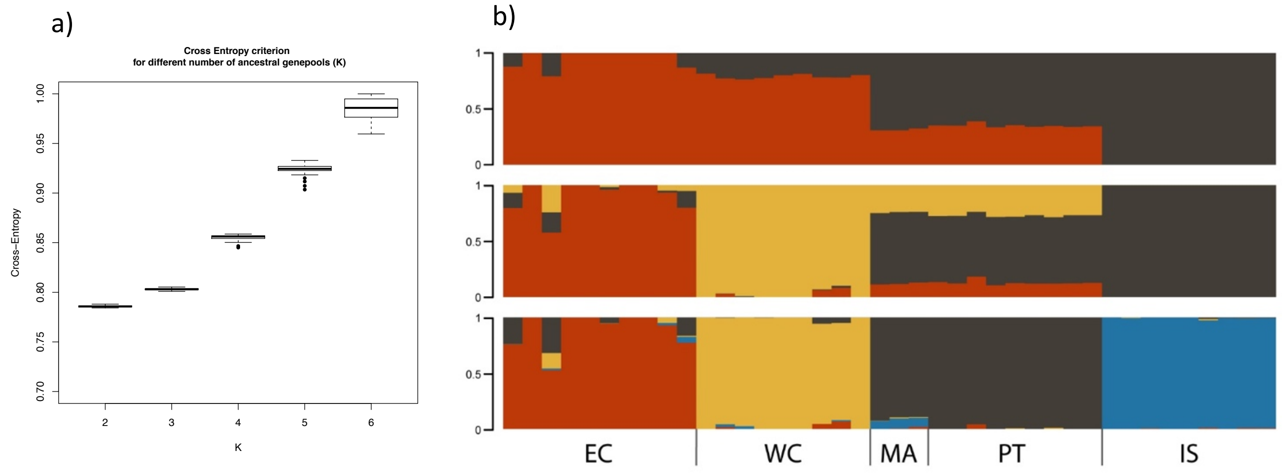

**Sup. Fig. 2** – Individual clustering estimated by sNMF. **a)** Values of the cross-entropy criterion for 25 sNMF runs per K, with K varying between 2 and 6. The lower values suggest a better fit for K=2. **b)** Individual ancestry estimated for K ranging between 2 to 4. Each vertical bar represents one individual, and the colours represent the relative contributions of each genetic lineage.

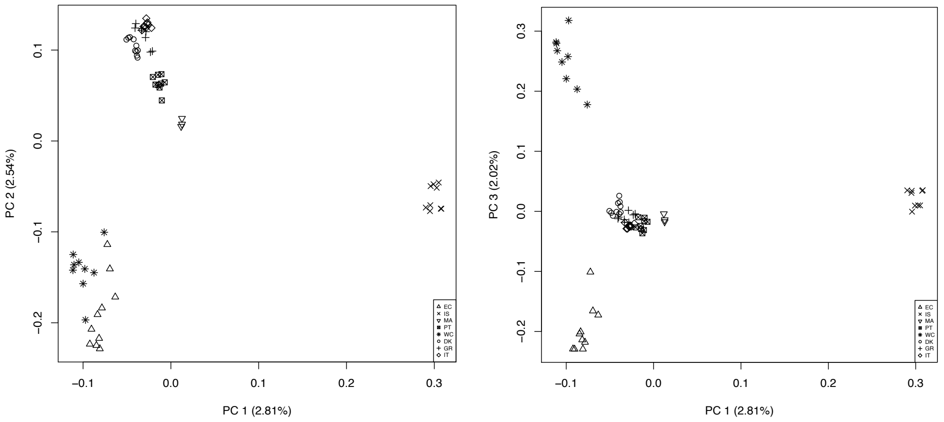

**Sup. Fig. 3** – PCA of individuals used in this study and European ones from Cumer et al. 2021. Axes 1 to 3 of the PCA are depicted.

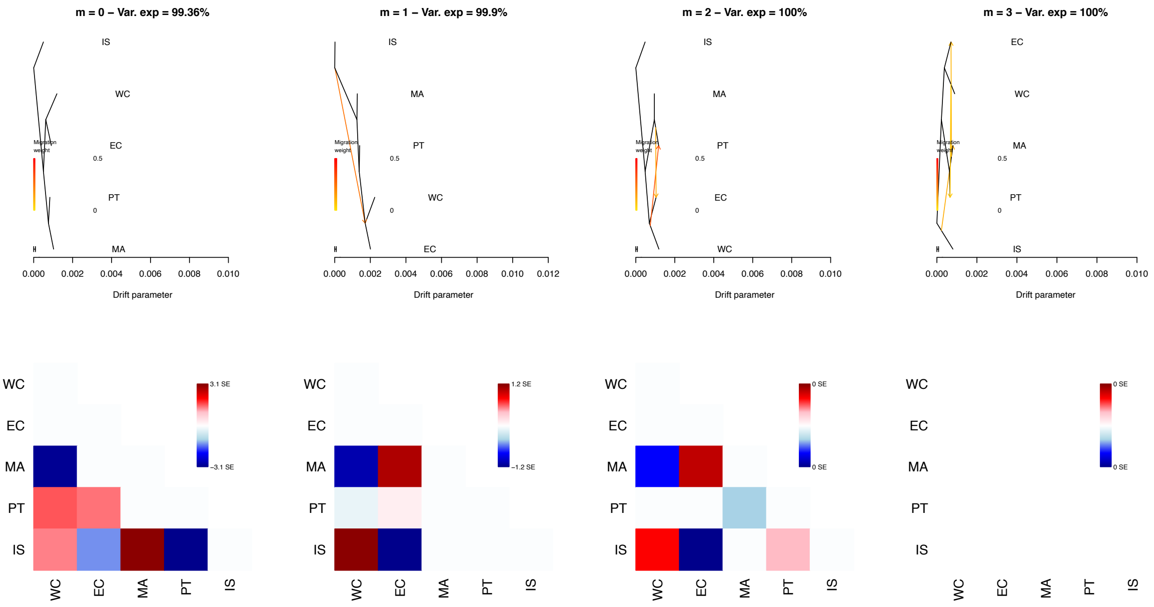

**Sup. Fig. 4** – Treemix analysis with 0 and up to 3 migration events (m), the variance explained, and their residual matrices. The best runs of each migration are depicted. Note that for 2 & 3 migration events, the residual error is 0 and the variance explained 100%; treemix was thus unable to add more than one migration events to the tree.

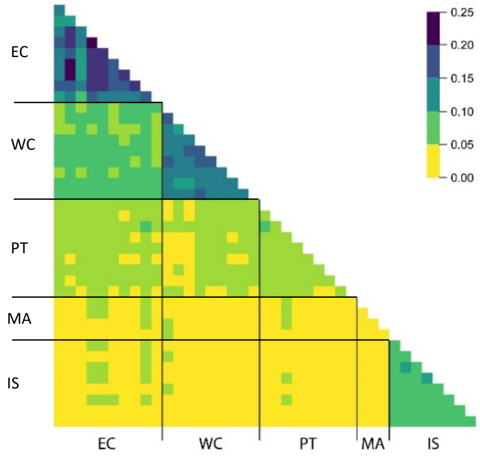

**Sup. Fig. 5** – Pairwise individual relatedness (β) heatmap between all individuals.

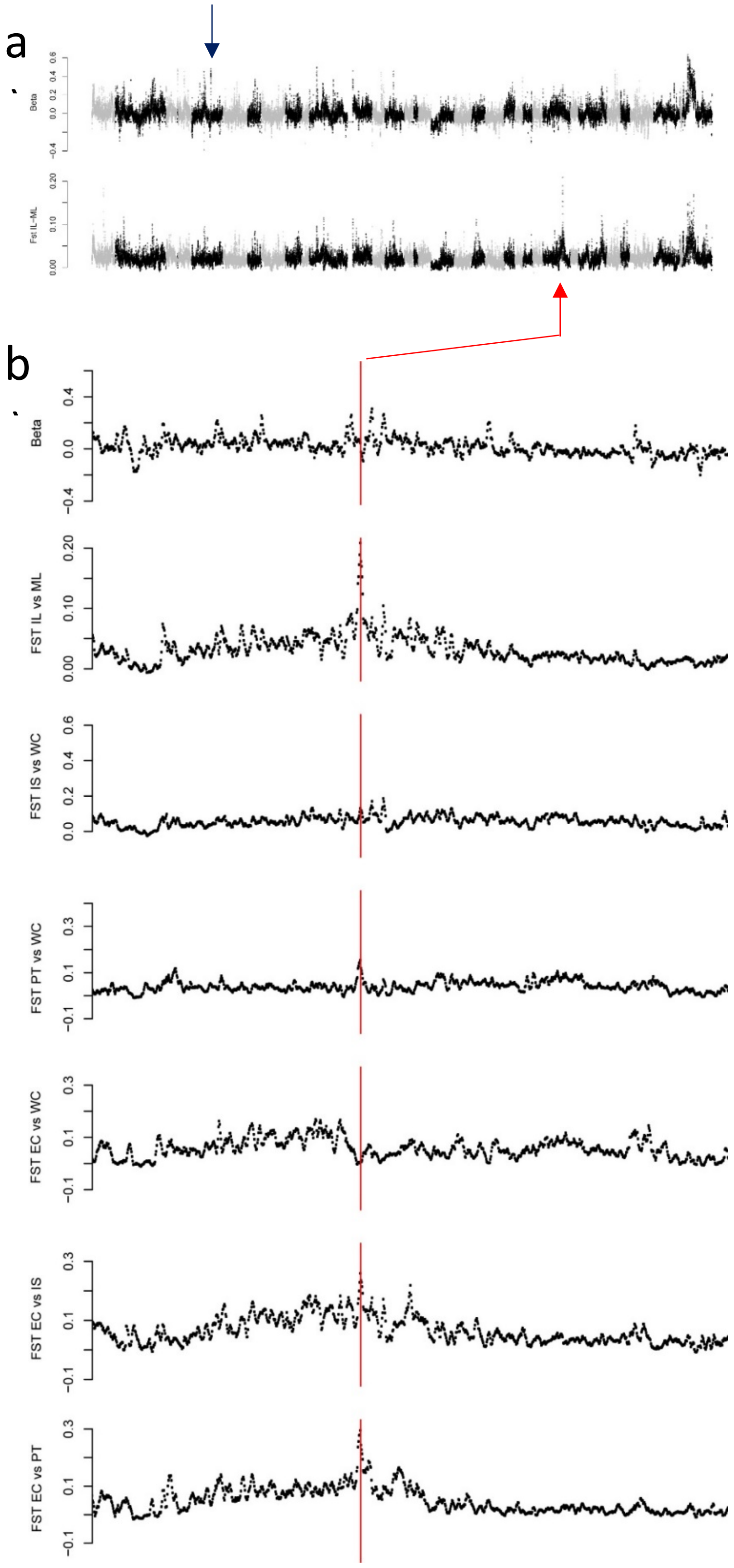

**Sup. Fig. 6** – Comparison of $F_{ST}^{Can}$ and $F_{ST}$ scans for **a)** the whole genome and **b)** a zoom on the highest $F_{ST}^{ILvsML}$ peak, absent from the $F_{ST}^{Can}$ scan (indicated in red). Note that this peak is actually a specificity of EC and not an overall difference between insular and mainland birds. $F_{ST}^{Can}$ did not produce such a peak, instead identifying regions that were actually common to both islands in the comparisons to the mainland (example indicated in blue).

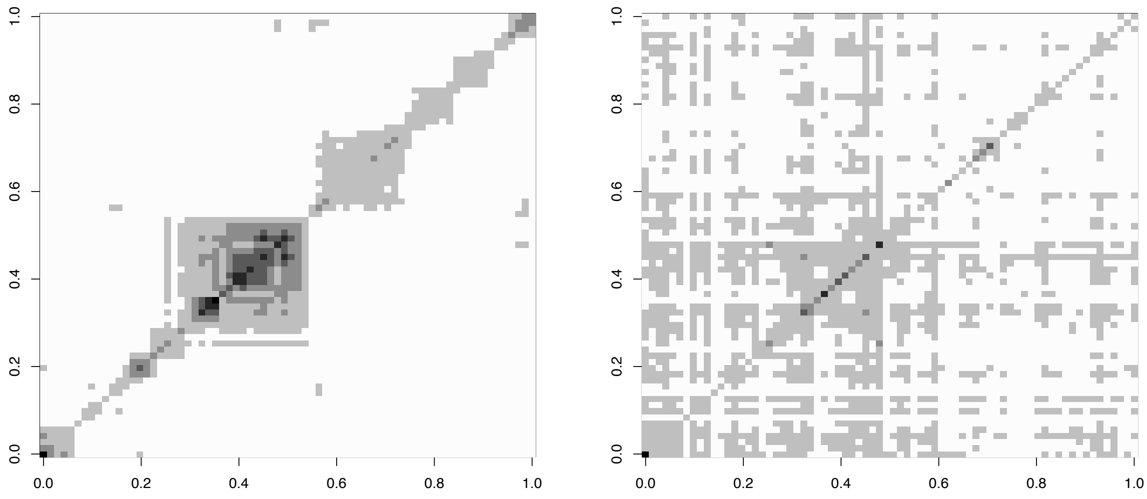

**Sup. Fig. 7** – Matrices of pairwise Linkage Disequilibrium (LD; $r^2$) along Scaffold 10006 of barn owls from Islands and from Mainland. Each pixel shows the average $r^2$ of 100 SNP, with higher values being of darker colour. The Island matrix clearly shows a region of amplified LD that the Mainland does not. This region corresponds to the highly differentiated segment shown in Figure 2.

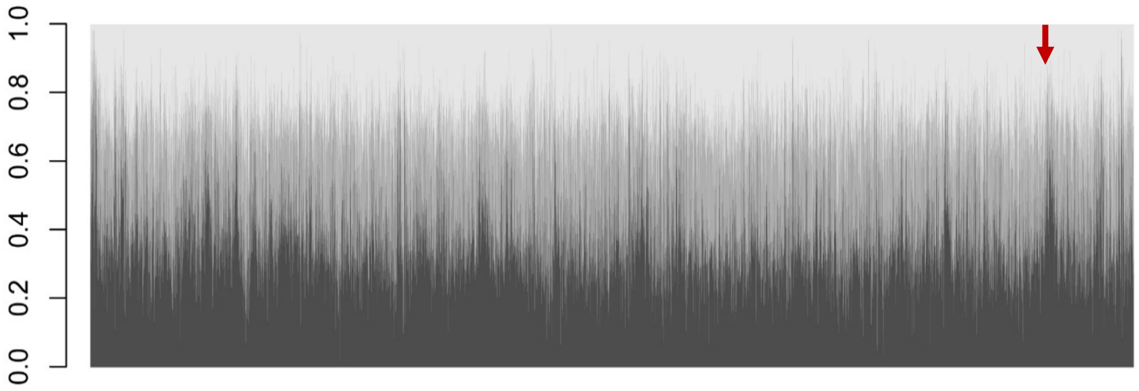

**Sup. Fig. 8** – Twisst tree weighting output along the genome. At each 100kb window, the proportion explained by each tree topology is shown in shades of grey (dark grey: ((WC,EC),PT,IS); grey: ((WC,PT),EC,IS); light grey: ((WC,IS),EC,PT)). Red arrow indicates the haplotype-like region selected on the islands (Figure 2). Note the increased proportion of dark grey trees, in which both island populations are placed in the terminal monophyletic branch, showing increased convergence among insular individuals in this genomic region.

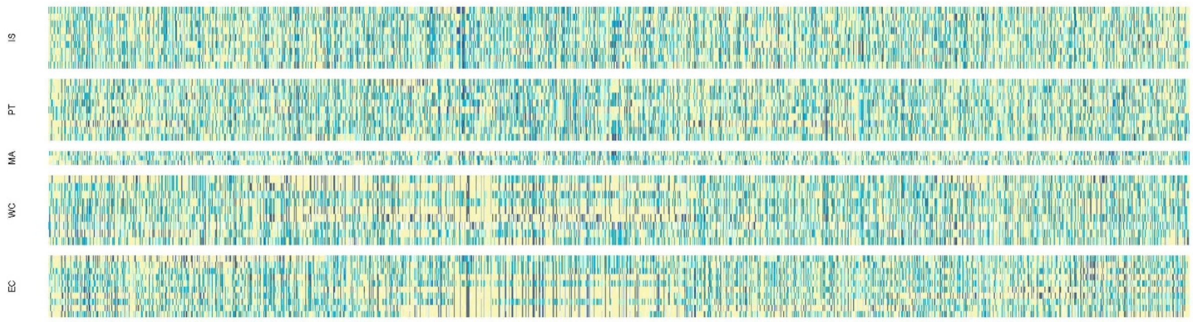

**Sup. Fig. 9** – Allele dosage representation along Scaffold 10006 of barn owls from the Canary Islands and the Mediterranean Basin. Each vertical bar represents the genotype at one biallelic SNP. In yellow, homozygote for one allele; light blue, heterozygote; dark blue, homozygote for the other allele. The highly differentiated region shown in Figure 2 is visible with in the EC and to a lesser extend WC populations but not in the others, with clear long yellow segments.

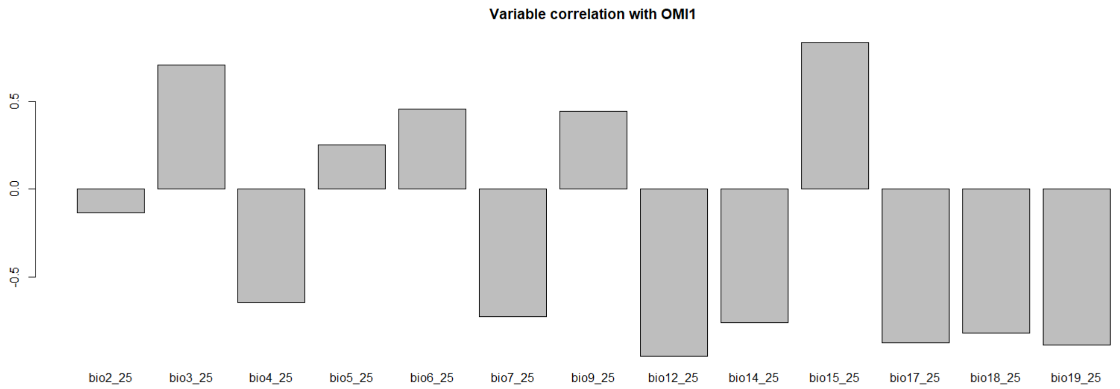

**Sup. Fig. 10** – Correlation of bioclim climatic variables with OMI1, the first axis of niche variance.
